# Revealing RNA virus diversity and evolution in unicellular algae transcriptomes

**DOI:** 10.1101/2021.05.09.443335

**Authors:** Justine Charon, Shauna Murray, Edward C. Holmes

## Abstract

Remarkably little is known about the diversity and evolution of RNA viruses in unicellular eukaryotes. We screened a total of 570 transcriptomes from the Marine Microbial Eukaryote Transcriptome Sequencing Project (MMETSP) project that encompasses a wide diversity of microbial eukaryotes, including most major photosynthetic lineages (i.e. the microalgae). From this, we identified 30 new and divergent RNA virus species, occupying a range of phylogenetic positions within the overall diversity of RNA viruses. Approximately one-third of the newly described viruses comprised single-stranded positive-sense RNA viruses from the order *Lenarviricota* associated with fungi, plants and protists, while another third were related to the order *Ghabrivirales*, including members of the protist and fungi-associated *Totiviridae*. Other viral species showed sequence similarity to positive-sense RNA viruses from the algae-associated *Marnaviridae*, the double-stranded RNA *Partitiviridae*, as well as a single negative-sense RNA virus related to the *Qinviridae.* Importantly, we were able to identify divergent RNA viruses from distant host taxa, revealing the ancestry of these viral families and greatly extending our knowledge of the RNA viromes of microalgal cultures. Both the limited number of viruses detected per sample and the low sequence identity to known RNA viruses imply that additional microalgal viruses exist that could not be detected at the current sequencing depth or were too divergent to be identified using sequence similarity. Together, these results highlight the need for further investigation of algal-associated RNA viruses as well as the development of new tools to identify RNA viruses that exhibit very high levels of sequence divergence.

## 1. Introduction

Viruses likely infect most, if not all, cellular species. For example, metagenomic studies of marine environments have revealed an enormous abundance and diversity of both DNA and RNA viruses (up to 10^8^ viruses/ ml)^1^ as well as their key role in biogeochemical processes^2^. Such ubiquity highlights the importance of obtaining a comprehensive picture of global virus diversity, including in host taxa that have only been poorly sampled to date^3^. Viruses of protists are a major exemplar of this untapped diversity.

Protists, defined as eukaryotic organisms that are not animal, plant, or fungi^4^, are highly diverse and include the algae. Some protists play a critical role in ecosystems as primary producers as well as being involved in nutrient cycling. Next generation sequencing (NGS) of protists has shown that their diversity is far greater than previously thought, with species numbers likely exceeding one million, although only a tiny fraction have been described to date^5^. In addition, protists have already proven to be an important source of virus diversity, with the giant *Mimiviridae* from the Amoebozoa a notable case in point^6^. Despite this, protist viruses remain largely overlooked, especially those associated with the unicellular microalgae. This is particularly striking in the case of RNA viruses: although RNA viruses were first described in unicellular algae in 2003^7^, they still comprise only 73 species from a very small number of algal lineages (Figure 1A)^8^.

**Figure 1.**
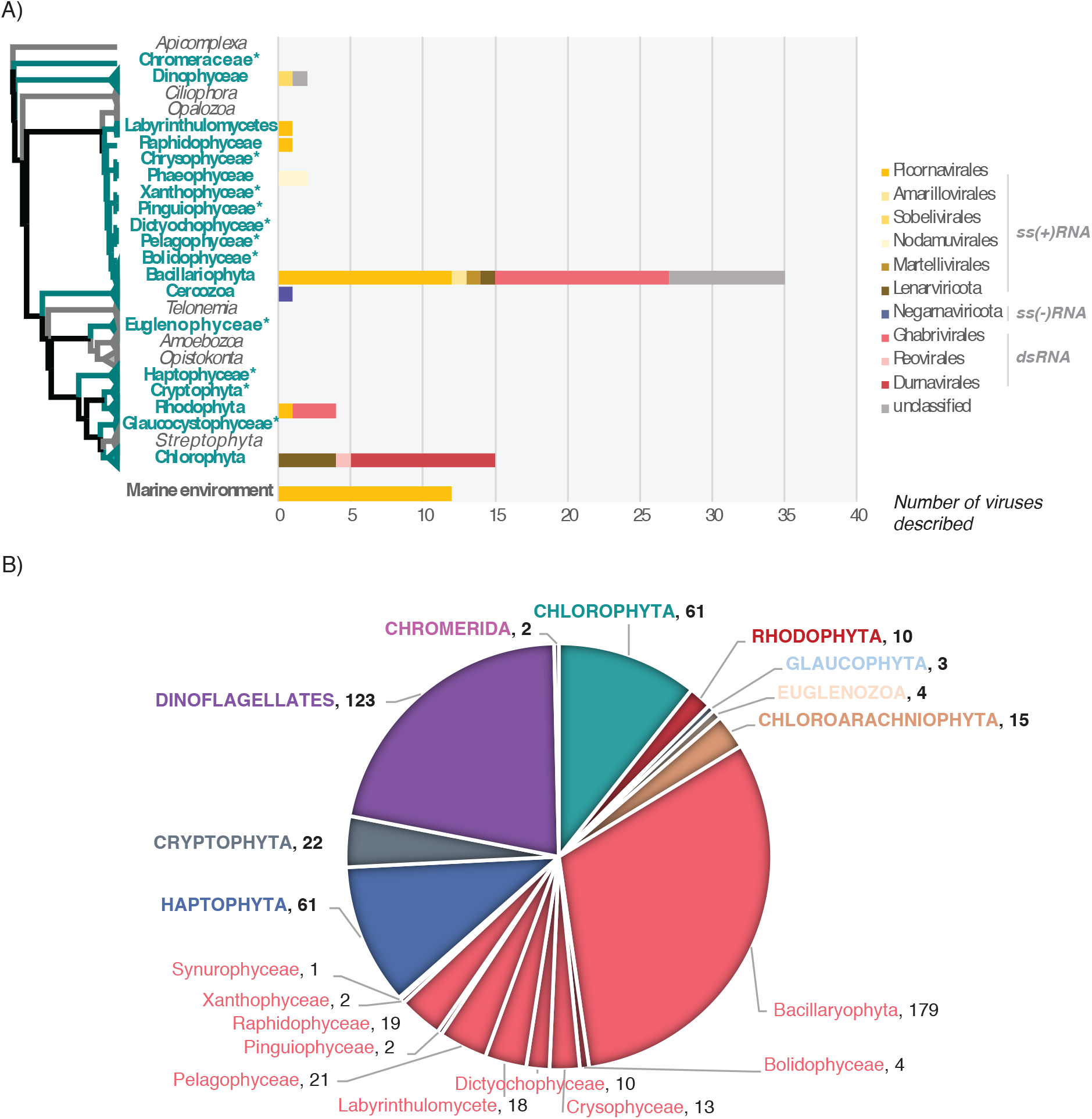
Currently reported RNA virus diversity in microalgae and the taxa studied here. **(A)** Left, Eukaryote phylogeny. The microalgae-containing eukaryotic lineages investigated here are highlighted in bold green. *Microalgae lineages for which no RNA viruses have been reported to date. Right, number of total viruses formally or likely associated with microalgae reported at NCBI (https://www.ncbi.nlm.nih.gov/labs/virus/vssi/), VirusHostdb (https://www.genome.jp/virushostdb/) and the literature. Viruses are coloured based on their taxonomy and genome composition. (**B)** Representative taxa from major algal lineages used in this study and the total number of transcriptomes analysed for each lineage.

There have been several metagenomic studies of viruses in aquatic microbial eukaryotes^9,10^. These have identified many thousands of virus sequences, with at least half predicted to have RNA genomes^11,12^. Similarly, metagenomics is proving a valuable means to mine viral diversity in uncultivable organisms^13^. However, because these studies have been conducted with environmental samples they cannot identify the specific host taxon with certainty.

This illustrates the inference gap between broad scale metagenomic surveys that identify huge numbers of new viral sequences, creating a large but unassigned depiction of the virosphere, and those studies based on virus isolation and detailed particle characterization, including cell culture, that are conducted on a very limited of number of viruses and create a highly accurate, but very narrow, vision of the virosphere^14^. However, establishing strong links between viruses and their specific hosts provides a firmer understanding of virus ecology and evolution, as well as virus-host interactions. Hence, the NGS-based investigation of RNA virus diversity from individual host species serves as a good compromise to fill the gap between large-scale virus detection through metagenomics and the detailed assignment of hosts through virus isolation and cell culture.

To better understand diversity of RNA viruses associated with microalgae, we performed viral metatranscriptomic analyses of data obtained from the Marine Microbial Eukaryote Transcriptome Sequencing Project (MMETSP)^15^. With 210 unique genera covering most unicellular algal-comprising lineages, the MMETSP constitutes the largest collection of transcriptome data collected from microbial eukaryote cultures, including axenic ones, and hence depicts a large component of eukaryotic diversity^15^ (Figure 1). Accordingly, we used both sequence and structural-based approaches to screen 570 transcriptomes from 19 major microalgae-containing lineages for the most conserved “hallmark” protein of RNA viruses – the RNA-dependent RNA polymerase (RdRp). To the best of our knowledge, this is the broadest exploration of RNA viruses conducted at the single host species level in microbial eukaryotes and the first attempt to identify RNA viruses in most of the microalgal lineages investigated here (Figure 1).

## 2. Methods

### 2.1 MMETSP contig retrieval

In total, 570 MMETSP accessions, corresponding to the microalgal-containing lineages, were included in this study. Contig data sets corresponding to each accession were retrieved from a Trinity re-assembly performed on the RNA-Seq data sets from MMETSP and available at https://doi.org/10.5281/zenodo.740440^16^. A description of all the transcriptome accessions and samples analysed here is available in Table S1.

### 2.2 ORF annotation

To optimize our computational analysis of the 570 contig data sets, we focused on those predicted to encode ORFs with a minimum length of 200 amino acids (assuming that shorter contigs would be too short to be included in a robust phylogenetic analyses). Accordingly, ORFs >200 amino acids in length were predicted using the GetORF tool from the EMBOSS package (v6.6.0). ORFs were predicted using the standard genetic code (with alternative initiation codons) as alternative genetic codes are not used in the microalgae analysed here^17^. The option -find 0 (translation of regions between STOP codons) was used to enable the detection of partial genomes, in which START codons could be missing due to partial virus genome recovery.

### 2.3 RNA virus sequence detection using sequence similarity

All predicted ORFs were compared to the entire non-redundant protein database (nr) (release April 2020) using DIAMOND BLASTp (v0.9.32)^18^ with the following options: --max-target-seqs 1 (top hit with best score retained) and an e-value cut-off of 1e-03. Additional sequence comparisons with identical BLASTp parameters were performed using either the newly-detected RdRp sequences or the RdRps from a previous large-scale analysis^12^ (available at ftp://ftp.ncbi.nih.gov/pub/wolf/_suppl/yangshan/rdrp.ya.fa).

To limit false-negative detection due to a bias in ORF prediction (in particular, partial genomes may not be detected due to their short length), all the contig nucleotide sequences were submitted to a RdRp protein database using DIAMOND BLASTx (v0.9.32, more sensitive option and 1e-03 e-value cut-off)^18^ to identify any additional RNA viruses. Top hits were retained and re-submitted against the entire nr protein database (April 2020 release) to remove false-positive hits (queries with a greater match to non-viral hits). All sequences retained from both the BLASTp and RdRp BLASTx analysis were manually checked to remove non-RNA virus sequences based on their taxonomy (predicted using the TaxonKit tool from NCBI; https://github.com/shenwei356/taxonkit).

All RNA virus-like sequences detected were functionally annotated using InterProscan (v5.39-77.0, default parameters) and non-RdRp sequences were filtered out. One sequence, sharing homology with the QDH87844.1 hypothetical protein H3RhizoLitter144407_000001, partial [Mitovirus sp.], was observed in 86 of the 570 data sets, including multiple species from multiple sampling locations. Considering the prevalence of this hit and the 100% identity between samples, we assumed this originates from environmental or sequencing-associated contamination. In addition, a small number of RNA virus-like sequences were identified based on their similarity to the RdRp from bovine viral diarrhea viruses 1 and 2 and considered biological product contaminants^19^. These were also discarded.

### 2.4 RNA virus sequence detection using protein profiles and 3D structures

In an attempt to detect more divergent viral RdRps we compared all the “orphan” ORFs (i.e. ORFs without any BLASTp hits at the 1e-03 e-value cut-off) against the viral RdRp-related profiles from the PFAM^20^ and PROSITE databases (Table S2) using the HMMer3 program^21^ (v3.3, default parameters, e-value<1e-05). An additional attempt to annotate orphan translated-ORFs was performed on the remaining sequences using the InterProscan software package from EMBL-EBI (v5.39-77.0, default parameters) (https://github.com/ebi-pf-team/interproscan).

The RdRp-like candidates identified in both the HMMer3 and InterProscan analysis were submitted to the Protein Homology/analogY Recognition Engine v 2.0 (Phyre2) web portal^22^ to confirm the presence of a RdRp signature (Table S3). Non-viral proteins (i.e. non-viral Phyre2 hit >90% confidence) were discarded, as were sequences with low HMM (e-value >1e-03) and Phyre2 scores (confidence level > 90%). Sequences that matched either the HMM RdRp (>1e-05) and/or Phyre2 RdRp (>90% confidence) were retained for further characterization as potential RNA viruses. In total, 80 RdRp-like candidates were quality-assessed by coverage analysis and manual checked for the presence of the standard A, B and C catalytic viral RdRp sequence motifs^23^ using Geneious (v11.1.4)^24^. Only those displaying related RdRp-like motifs were retained as potential RdRp protein candidates (Table S3).

### 2.5 Contig manual extension and genome annotation

Full-length nucleotide sequences encoding the protein retained from the sequence-based and structure-based detection approaches were retrieved and used as references for mapping SRA reads corresponding to each sample (BioProject PRJNA231566) using the SRA extension package of Bowtie2 (v2.3.5.1-sra)^25^. Read coverages of each contig were checked using Geneious (v11.1.4) and, when needed, extremities were manually extended and contigs re-submitted to read mapping, until no overhanging extremities were observed.

The relative abundance of each putative viral sequence was reported as the number of reads per million: that is, the number of reads mapping to the contig divided by the total number of reads of the corresponding SRA library multiplied by one million. Poorly-represented viral sequences were considered as potential cross-library contaminants derived from index-hopping and discarded when they accounted for less than 0.1% of the highest abundance of the same sequence in another library^26^.

Genomic organizations were constructed using Geneious (v11.1.4). ORFs were predicted using the standard genetic code or, when suitable, using alternative mitochondrial or plastid-associated genetic codes. Tentative virus names were taken from Greek mythology.

### 2.6 Host rbcL gene abundance estimation

To estimate levels of virus abundance in comparison to those from their putative hosts, the abundance of the host Ribulose bisphosphate carboxylase large chain (*rbcL*) gene was assessed using the Bowtie2 SRA package (v2.3.5.1-sra) and mapped to SRA reads from the *rbcL* gene of each corresponding species (whenever available)^25^. The SRA and *rbcL* gene accessions used are reported in Table S4.

### 2.7 Secondary host profiling

According to the MMETSP sample requirements, all cultures were subjected to SSU rRNA sequencing to ensure they were mono-strain and not contaminated with additional microbial eukaryotes. Nevertheless, the presence of other microbial contaminants was possible. As we expect most of the potential Archaea and Bacteria contaminants will not have an available genome sequence, their profiling in the samples was performed by analysing the closest homologs of each contig using both BLASTn (BLAST+ package, v2.9.0) and BLASTp (DIAMOND, v2.0.4) against the nt and nr databases, respectively. Contigs were grouped at the kingdom level based on the taxonomic affiliation of their closest homologs in the databases, with the abundance of each kingdom defined as the sum of each contig abundance value (transcripts per million)^16^.

### 2.8 Phylogenetic analysis

For each virus phylum and order, the RefSeq and most closely related RdRp sequences were retrieved from GenBank and aligned with newly identified RdRp sequences using the L-INS-I algorithm in the MAFFT program (v7.402)^27^. Resulting sequence alignments were trimmed using TrimAl to remove ambiguously aligned regions with different levels of stringency, optimized for each alignment (v1.4.1, “automated1” mode). Maximum likelihood phylogenies based on amino acid alignments were inferred using IQ-TREE (v2.0-rc1)^28^, with ModelFinder used to find the best-fit substitution model in each case (see figure legends)^29^ and both the SH-like approximate likelihood ratio test and ultrafast nonparametric bootstrap (1000 replicates) used to assign support to individual nodes^30^. All phylogenies were visualized, and mid-point rooted (for clarity only) using the Figtree software (v1.4.4).

### 2.9 Detection of endogenous viral elements

To determine whether any of the newly detected viral sequences were endogenous viral elements (EVEs) rather than true exogenous viruses, the nucleotide sequences of viral candidates were used as a query for BLASTn (online version, default algorithm parameters) against corresponding host genome sequence, whenever available.

## 3. Results

### 3.1 Overall virus diversity

Our analysis of the 570 MMETSP transcriptomes obtained from 247 total microalgal species spread over 10 major groups of algae (Table 1B) identified 30 new RNA viral species. These newly identified viruses largely represented the single-stranded positive-sense RNA (ssRNA+) virus phylum *Lenarviricota* and the order *Picornavirales* (Figure 2A and B), as well as the double-stranded (dsRNA) RNA virus orders *Durnavirales* and *Ghabrivirales* (Figure 2C and D). A single negative-sense RNA (ss-RNA) virus was also identified in *Pseudo-nitzchia heimii* that fell within the *Qinviridae* (order *Muvirales*).

**Table 1.** List of new RNA viruses discovered in this study. Read abundances are indicated as the number of reads per million. Likely hosts correspond to eukaryotic lineages detected at levels using BLASTn/BLASTp analysis and phylogenies.

| Virus name | MMETSP sample<br>(Phylum/class) | Genome<br>status | Reads/<br>million | BLASTp best hits<br>(GenBank acc./Organism) | %ID | E-value | Likely host(s)<br>(BLAST) | Likely host(s)<br>(Phylogenies) | Proposed host |
| --- | --- | --- | --- | --- | --- | --- | --- | --- | --- |
| Amphitrite narna-like virus | MMETSP1061<br>*P. pungens*<br>(Bacillariophyta) | Full-length | 48 | QIR30281.1 RdRp<br>[Plasmopara viticola associated<br>narnavirus 2] | 41 | 5E-144 | Bacillariophyta | Fungi/Protist | Bacillariophyta |
| Poseidon narna-like virus | MMETSP0418<br>*A. radiata*<br>(Bacillariophyta) | Partial | 8 | QDH89392.1 RdRp, partial<br>[Mitovirus sp.] | 34 | 4E-17 | Bacillariophyta | Marine<br>arthropod | Bacillariophyta |
| Halia narna-like virus | MMETSP0418<br>*A. radiata*<br>(Bacillariophyta) | Full-length | 108 | QBC65281.1 RdRp, partial<br>[Rhizopus microsporus 23S<br>narnavirus] | 32 | 4E-17 | Bacillariophyta | Protist | Bacillariophyta |
| Triton levi-like virus | MMETSP1471<br>*P. provasolii*<br>(Chlorophyta) | Partial | 64 | APG76993.1 hypothetical<br>protein [Beihai levi-like virus<br>20] | 46 | 3E-65 | Chlorophyta;<br>Bacteria | Bacteria | Bacteria |
| Aiolos mito-like virus | MMETSP0286<br>*P. polylepis*<br>(Haptophyta) | Full-length | 54 | YP_009272901.1 RdRp<br>[Fusarium poae mitovirus 4] | 35 | 3E-38 | Haptophyta | Sea sponge | Haptophyta |
| Asopus mito-like virus | MMETSP0164<br>*C. braarudii*<br>(Haptophyta) | Partial | 12 | QDM55307.1 RdRp<br>[Geopora sumneriana mitovirus<br>1] | 34 | 2E-35 | Haptophyta | Sea sponge | Haptophyta |
| Athena mito-like virus | MMETSP0719<br>*C. curvisetus*<br>(Bacillariophyta) | Partial | 54 | ASM94070.1 putative RdRp,<br>partial [Barns Ness breadcrumb<br>sponge narna-like virus 5] | 65 | 6E-72 | Bacillariophyta;<br>Bacteria | Sea sponge | Bacillariophyta |
| Daimones mito-like virus | MMETSP0286 | Full-length | 104 | YP_009552787.1 RNA-directed<br>RNA polymerase | 26 | 4E-16 | Haptophyta | Freshwater<br>arthropods | Haptophyta |
|  | *P. polylepis*<br>(Haptophyta) |  |  | [Rhizophagus sp. RF1<br>mitovirus] |  |  |  |  |  |
| Despoena mito-like<br>virus | MMETSP0167<br>*R. maculata*<br>(Rhodophyta) | Full-length | 115 | ALM62241.1 RdRp<br>[Soybean leaf-associated<br>mitovirus 1] | 34 | 6E-32 | Rhodophyta;<br>Bacteria | Freshwater<br>arthropods | Rhodophyta |
| Proteus mito-like<br>virus | MMETSP1081<br>*P. amyliifera*<br>(Chlorophyta) | Full-length | 388 | ALM62242.1 RdRp<br>[Soybean leaf-associated<br>mitovirus 2] | 32 | 7E-46 | Chlorophyta | Fungi/Protist | Chlorophyta |
| Telchines mito-like<br>virus | MMETSP0725<br>*Amphiprora*<br>(Bacillariophyta) | Partial | 15 | QDA33961.1 RdRp<br>[Mitovirus 1 BEG47] | 25 | 5E-21 | Bacillariophyta | Algae | Bacillariophyta |
|  | MMETSP0724<br>*Amphiprora*<br>(Bacillariophyta) | Partial | 26 |  |  |  |  |  |  |
| Susy yue-like virus | MMETSP1423<br>*P. heimii*<br>(Bacillariophyta) | Partial | 5 | QDH86724.1 RdRp, partial<br>[Qinviridae sp.] | 42 | 1E-21 | Bacillariophyta | Soil samples/<br>Marine<br>arthropod | Bacillariophyta |
| Aethusa amalga-like virus | MMETSP0011<br>*R. marinus*<br>(Rhodophyta) | Partial | 83 | ANN12897.1 putative CP/RdRp<br>[Zygosaccharomyces bailii<br>virus Z] | 43 | 2E-12 | Rhodophyta;<br>Bacteria | Marine<br>arthropod | Rhodophyta |
| Benthesicyme<br>durna-like virus | MMETSP1319<br>*T. pacifica*<br>(Bolidophyceae) | Partial | 404 | QDH90748.1 RdRp, partial<br>[Partitiviridae sp.] | 29 | 1E-17 | Bolidophyceae | Protist | Bolidophyceae |
| Herophile durna-like virus | MMETSP0140<br>*P. australis*<br>(Bacillariophyta) | Partial | 10 | QOW97238.1 RdRp<br>[Amalga-like lacheneauvirus] | 27 | 2E-19 | Bacillariophyta | Chlorophyta | Bacillariophyta |
| Cymopoleia durna-like virus | MMETSP1081<br>*P. amyliifera*<br>(Chlorophyta) | Partial | 10 | YP_009551448.1 RdRp<br>[Diatom colony associated<br>dsRNA virus 2] | 31 | 2E-34 | Chlorophyta | Fungi | Chlorophyta |
| Ourea durna-like virus | MMETSP0797<br>*D. acuminata*<br>(Dinophyceae) | Partial | 4 | ARO72610.1 RdRp<br>[Spinach deltapartitivirus 1] | 27 | 4E-11 | Dinophyceae;<br>Bacteria | Land plant | Dinophyceae |
| Aegean partiti-like virus | MMETSP0491<br>*T. chuii*<br>(Chlorophyta) | Full-length | 3296 | QOW97235.1 RdRp<br>[Partiti-like lacotivirus] | 29 | 6E-62 | Chlorophyta | Chlorophyta | Chlorophyta |
| Pelias marna-like virus | MMETSP1377<br>*Symbiodinium sp.*<br>(Dinophyceae) | Full-length | 60553 | YP_009337401.1 hypothetical<br>protein 2<br>[Wenzhou picorna-like virus 4] | 26 | 8E-98 | Dinophyceae | Algae | Xanthophyceae |
| Neleus marna-like virus, 1 | MMETSP0946<br>*V. litorea*<br>(Xanthophyceae) | Full-length | 806763 | YP_009336927.1 hypothetical<br>protein 1<br>[Shahe picorna-like virus 3] | 33 | 3E-180 | Vaucheriaceae | Algae | Xanthophyceae |
| Neleus marna-like virus, 2 | MMETSP0945<br>*V. litorea*<br>(Xanthophyceae) | Full-length | 711119 | YP_009336927.1 hypothetical<br>protein 1<br>[Shahe picorna-like virus 3] | 33 | 4E-180 | Vaucheriaceae | Algae | Xanthophyceae |
| Tyro marna-like virus | MMETSP0905<br>*T. antarctica*<br>(Bacillariophyta) | Partial | 126 | YP_001429582.1 hypothetical<br>protein JP-A_gp2<br>[Marine RNA virus JP-A] | 75 | 3E-272 | Bacillariophyta;<br>Bacteria | Algae | Bacillariophyta |
|  | MMETSP0903<br>*T. antarctica*<br>(Bacillariophyta) | Partial | 2034 |  |  |  |  |  |  |
|  | MMETSP0902<br>*T. antarctica*<br>(Bacillariophyta) | Partial | 237 |  |  |  |  |  |  |
| Aloadae toti-like virus,1 | MMETSP1388<br>*Isochrysis*<br>(Haptophyta) | Partial | 39 | QIJ70132.1 RdRp<br>[Keenan toti-like virus] | 33 | 2E-109 | Haptophyta | Fungi<br>/Invertebrates | Haptophyta |
| Aloadae toti-like virus,2 | MMETSP1090 | Partial | 11 | QIJ70132.1 RdRp<br>[Keenan toti-like virus] | 33 | 2E-109 | Haptophyta | Fungi<br>/Invertebrates | Haptophyta |

|  | *Isochrysis*<br>(Haptophyta) |  |  |  |  |  |  |  |  |
| --- | --- | --- | --- | --- | --- | --- | --- | --- | --- |
| Antaeus toti-like virus,1 | MMETSP0154<br>*T. antarctica*<br>(Bacillariophyta) | Full-length | 27 | QGY72637.1 putative coat protein [Plasmopara viticola associated totivirus-like 2] | 22 | 1E-10 | Bacillariophyta | Protist | Bacillariophyta |
| Antaeus toti-like virus,2 | MMETSP0152<br>*T. antarctica*<br>(Bacillariophyta) | Full-length | 7 | BBJ21451.1 CP-RdRp fusion protein [Pythium splendens RNA virus 1] | 40 | 5E-53 | Bacillariophyta | Protist | Bacillariophyta |
| Charybdis toti-like virus | MMETSP0853<br>*P. fraudulenta*<br>(Bacillariophyta) | Partial | 38 | YP_003288763.1 RdRp [Rosellinia necatrix megabirnavirus 1/W779] | 30 | 4E-24 | Bacillariophyta; Bacteria | Fungi | Bacillariophyta |
|  | MMETSP0851<br>*P. fraudulenta*<br>(Bacillariophyta) | Partial | 44 |  |  |  |  |  |  |
|  | MMETSP0850<br>*P. fraudulenta*<br>(Bacillariophyta) | Partial | 41 |  |  |  |  |  |  |
|  | MMETSP0852<br>*P. fraudulenta*<br>(Bacillariophyta) | Partial | 14 |  |  |  |  |  |  |
| Chrysaor toti-like virus | MMETSP0418<br>*A. radiata*<br>(Bacillariophyta) | Partial | 40 | YP_009551502.1 RdRp [Diatom colony associated dsRNA virus 17 genome type B] | 27 | 9E-95 | Bacillariophyta; Bacteria | Soil | Bacillariophyta |
| Laestrygon toti-like virus | MMETSP1451<br>*V. brassicaformis*<br>(Chromeraceae) | Partial | 29 | YP_009551504.1 RdRp [Diatom colony associated dsRNA virus 17 genome type A] | 34 | 4E-112 | Chromeraceae | Soil | Chromeraceae |
| Arion toti-like virus | MMETSP0796<br>*P. bahamense*<br>(Dinophyceae) | Partial | 31 | QGA70930.1 RdRp<br>[Ahus virus] | 25 | 3E-18 | Dinophyceae;<br>Bacteria | Protist/<br>Marine host | Dinophyceae |
| Otus toti-like virus | MMETSP0011<br>*R. marinus*<br>(Rhodophyta) | Full-length | 31 | AMB17469.1 RdRp, partial<br>[Delisea pulchra totivirus IndA] | 51 | 3E-120 | Rhodophyta;<br>Bacteria | Fungi | Rhodophyta |
| Polyphemus toti-like virus | MMETSP0418<br>*A. radiata*<br>(Bacillariophyta) | Partial | 10 | YP_009552789.1 RdRp<br>[Diatom colony associated<br>dsRNA virus 5] | 59 | 3E-79 | Bacillariophyta;<br>Bacteria | Algae/<br>Protist | Bacillariophyta |
| Ephialtes toti-like virus | MMETSP0418<br>*A. radiata*<br>(Bacillariophyta) | Partial | 13 | YP_009552789.1 RdRp<br>[Diatom colony associated<br>dsRNA virus 5] | 63 | 1E-200 | Bacillariophyta;<br>Bacteria | Algae/<br>Protist | Bacillariophyta |

**Figure 2.**
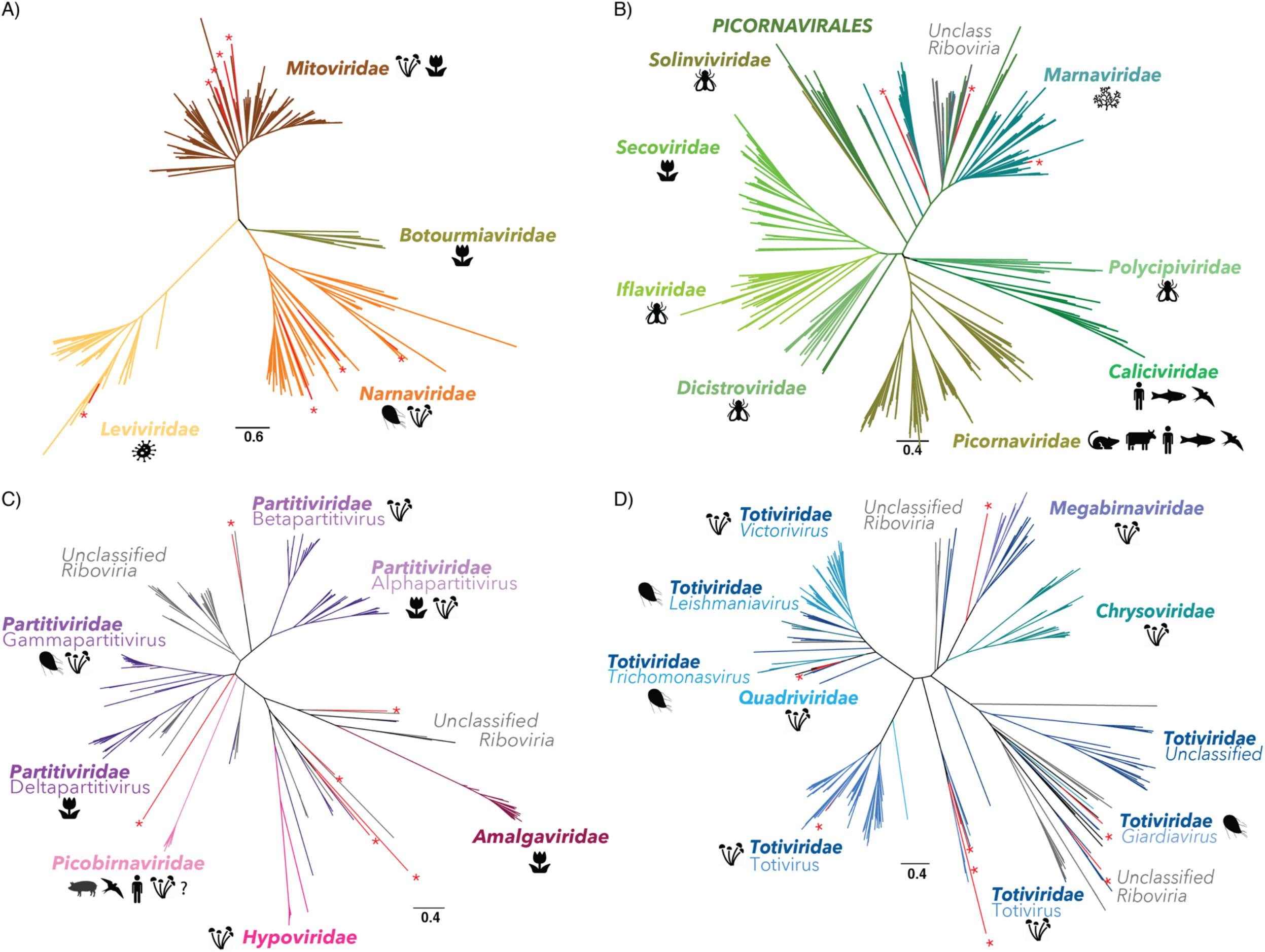
Newly described RNA virus sequences within the diversity of RNA viruses using RdRp phylogenies. Newly described sequences are indicated in red with “*” symbols. Phylogenies of: (A) the phylum *Lenarnaviricota* (ssRNA+); (B) the order *Picornavirales* (ssRNA+); (C) the order *Durnavirales* (dsRNA); (D) the order *Ghabrivirales* (dsRNA). For each viral family, the host range was retrieved from VirusHostdb and the ICTV report^31,32^.

Notably, all the RdRps identified in the BLAST analysis exhibited very high levels of sequence divergence, with median pairwise identity values of only ~35% to the closest known virus homolog (Table 1). In addition, with the exceptions of Pelias marna-like virus and Neleus marna-like virus, the newly described viral sequences were at relatively low abundance all (Table 1). This may reflect the lack of an rRNA depletion step used in the MMETSP library preparation, such that any RNA viruses would necessarily only represent a small proportion of reads. To shed more light on this issue, we compared levels of virus abundance with the expression levels of a host gene – that encoding the large subunit of the Ribulose-1,5-Bisphosphate Carboxylase/Oxygenase *(rbcL*) (Figure S1, Table S4). The *rbcL* gene is commonly used as a diversity marker in algae^33^, and sequences are available for all the microalgal species used here. Overall, the number of reads mapping to putative RNA viruses are in the same order of magnitude or higher than those reported for the host *rbcL* gene (Figure S1), compatible with their designation as replicating viruses.

### 3.2 Additional cellular organisms in the transcriptome data

We used mono-strain cultures of microbial eukaryotes to investigate the relationship among RNA viruses and their hosts. While the lack of additional eukaryotic organisms (fungi, other protists) was supposedly ensured under the MMETSP project guidelines, with 18S rRNA sequencing of each culture^15^, some caveats remain for non-axenic cultures (Table S5). Indeed, some cultures likely contain contaminating Bacteria or Archaea, sometimes as intracellular parasites or as obligate mutualists in the culture media ^5^. To assess this, contigs from libraries positive for RNA viruses were submitted to BLASTn and BLASTx. The ratio of assigned contigs and their kingdom assignments are summarized Figure 3 and used to infer the likely host organisms (Table 1).

**Figure 3.**
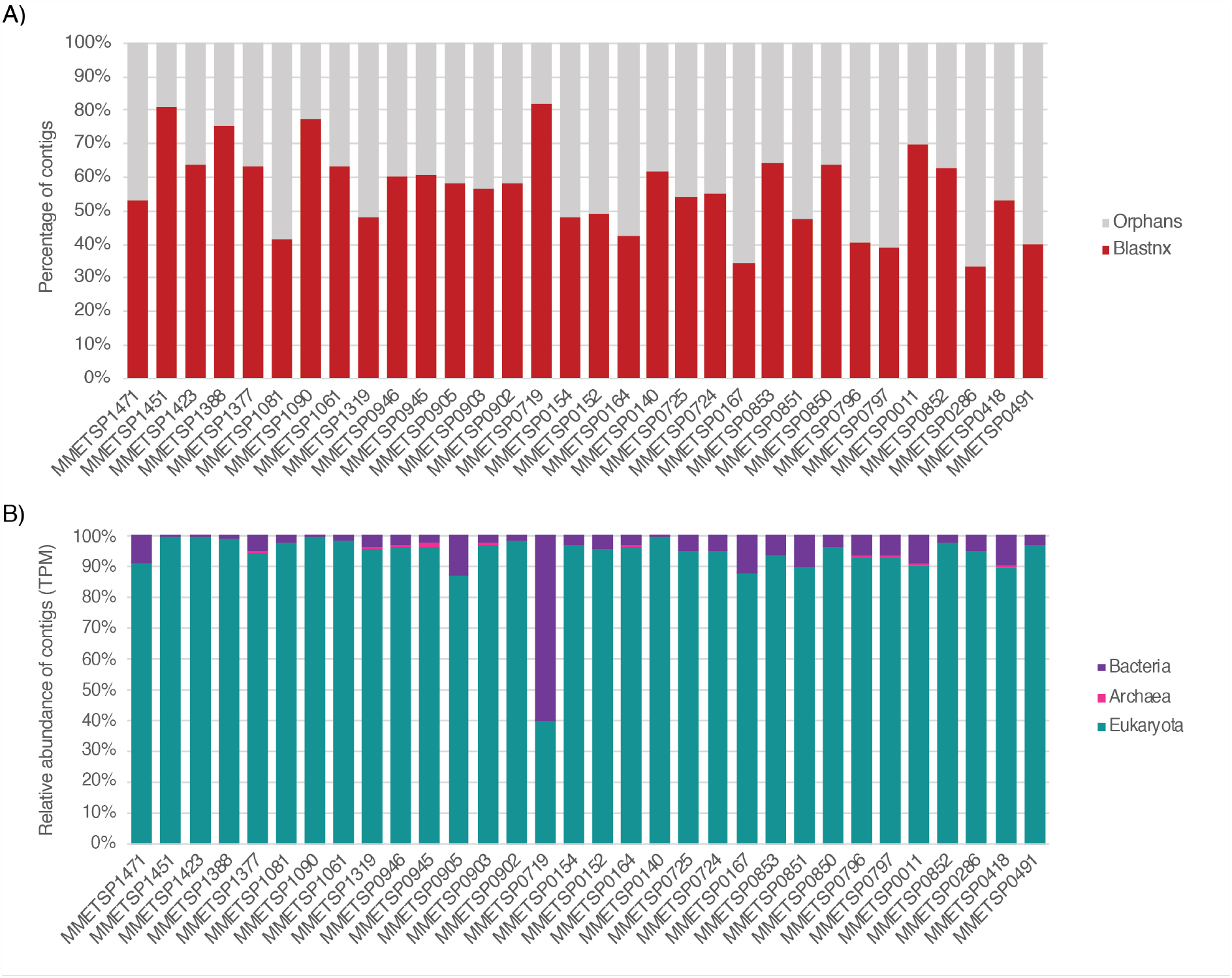
Taxonomic assignment of contigs in RNA virus positive MMETSP libraries. (A) Ratio of contigs with hits to the nt and nr databases (red) versus orphans contigs (grey). (B) Relative abundance of cellular organism-like contigs based on the taxonomic assignment of their closest homologs in the nr and nt databases at the kingdom level. Contig abundances are calculated as transcripts per million (TPM).

Approximately half of the total contigs identified here could not be assigned using BLAST approaches (Figure 3A), with prokaryotic organisms on average representing less than 10% of assigned contigs (Figure 3B). However, the MMETSP0719 containing *C. curvisetus* (Bacillariophyta) is enriched with co-infecting bacteria. According to the BLASTn/BLASTp entries obtained for this sample, this seems largely due to the presence of the marine alphaproteobacteria *Jannaschia*. This is to be expected as some algal species require the presence of particular bacterial species to obtain essential nutrients ^34^.

### 3.3 Distribution and prevalence of RNA viruses in MMETSP cultured strains

We found evidence for RNA viruses – that is, hits to the viral RdRp – in eight of the 19 major groups of microalgae, without detectable virus/algal taxon specificity (Figure 4).

**Figure 4.**
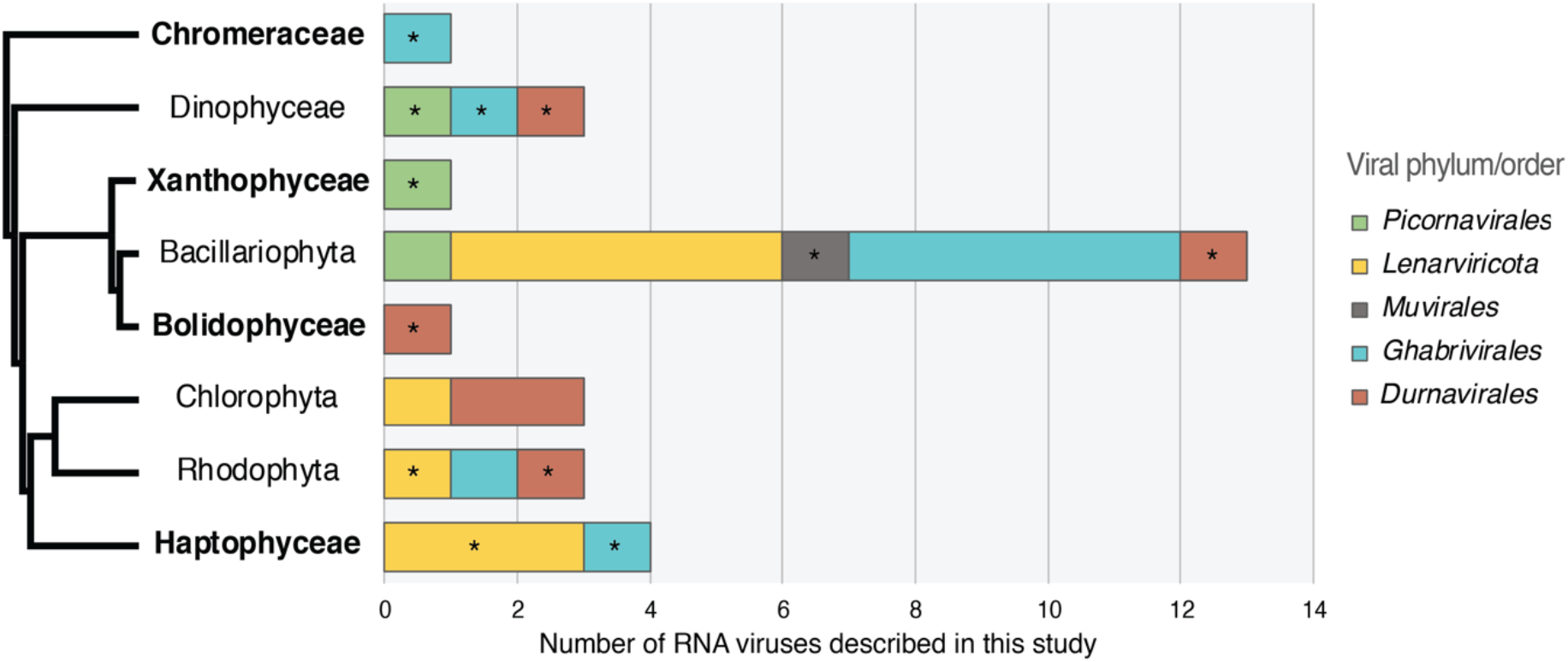
Distribution of RNA virus groups identified in algae. Only algal lineages containing RNA virus RdRps are shown. Left, cladogram of the algal host lineages positive for RNA viruses. Taxa for which no RNA viruses have previously been reported are indicated in bold. Right, total counts of newly described RNA viral sequences in each algal taxon (including viruses observed in several samples from the same taxa). *First observation of this virus taxon in the corresponding algal clade. The levi-like sequence that likely infects a bacterial host was excluded.

The distribution of RNA viruses is highly heterogeneous among the microalgae studied here, with a large representation in the Bacillariophyta, Dinophyceae and Haptophyceae, with only few or no viruses in the other taxa analyzed here (Figure 4). It is important to note that the number of viruses is strongly associated with the number of libraries analysed and thus likely depicts a limit of detection imposed by small sample sizes in some groups (i.e. large numbers of transcriptomes are available for the Bacillariophyta, Dinophyceae and Haptophyceae).

### 3.4 Positive-sense RNA viruses (ssRNA+)

Eleven of the 30 viruses discovered in this study show clear homology to three of the four families that comprise the recently classified phylum *Lenarviricota* of ssRNA+ viruses: the *Leviviridae*, the *Narnaviridae* and the *Mitoviridae* (Table 1). In all cases, levels of RdRp identities to the closest homologs were <60%, reflecting high levels of sequence divergence and leading us to propose that these 11 sequences are novel viral species (**Table 1**).

#### 3.4.1 *Narnaviridae*-like sequences

Three RdRp-containing contigs – denoted Amphitrite narna-like virus, Poseidon narna-like virus and Halia narna-like virus – were related to the *Narnaviridae*, occupying diverse positions in a phylogeny of this virus family (Figure 2 and Figure 5).

**Figure 5.**
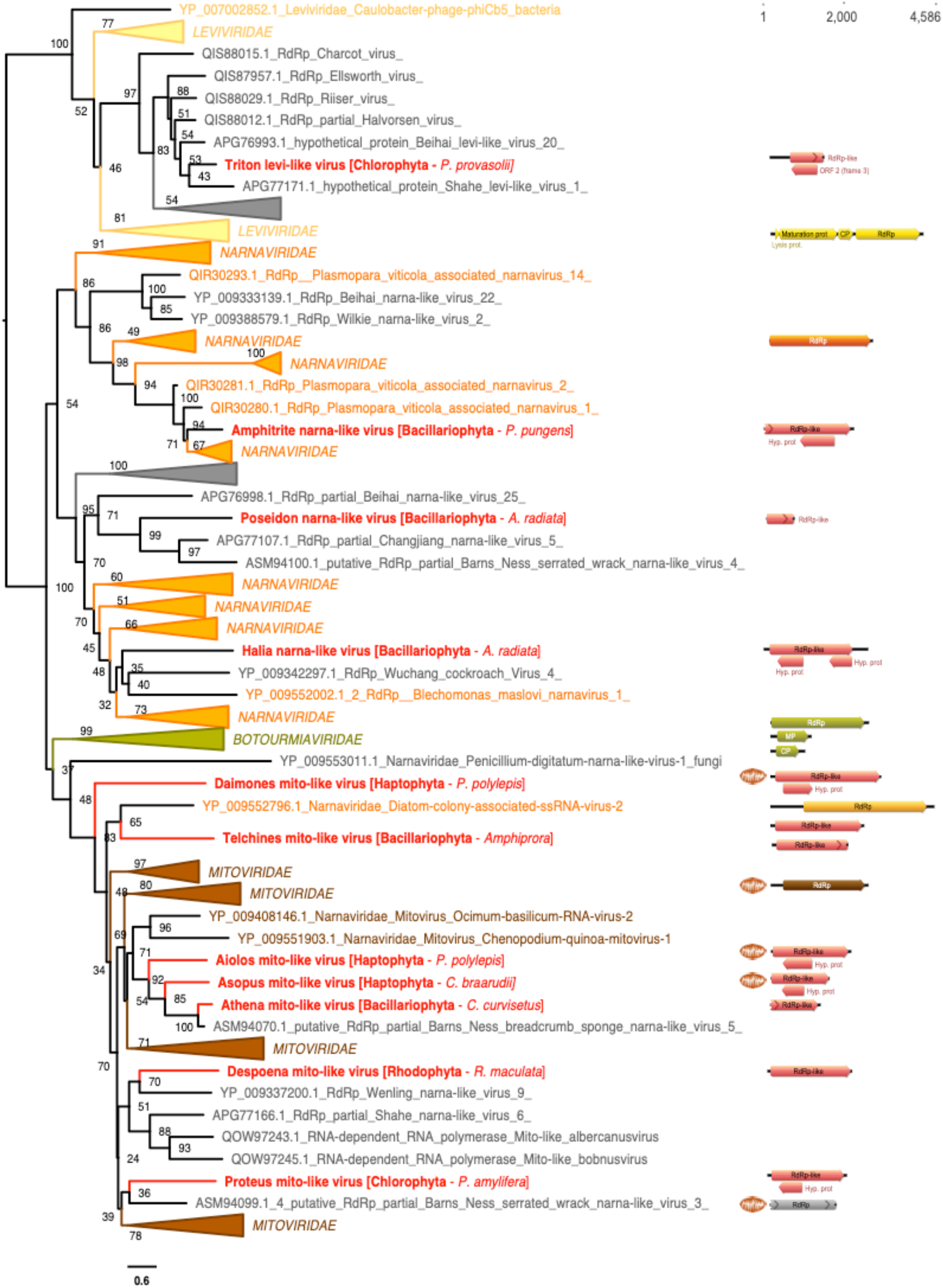
Phylogenetic position of the newly described RNA virus sequences in the phylum *Lenarviricota*. Left: ML phylogeny of the *Lenaviricota* RdRp (LG+F+R8 amino acid substitution model). Newly described viruses are shown in red. Algal host taxa are specified in brackets. Branch labels = bootstrap support (%). The tree is mid-point rooted for clarity only. Right: genomic organisation of the newly described viruses (red), closest homologs and *Lenarviricota* RefSeq representatives: Cassava virus C (NC_013111; *Botourmiaviridae*), Saccharomyces 23S RNA (NC_004050; *Narnaviridae*), Acinetobacter phage AP205 (NC_002700; *Leviviridae*), Chenopodium quinoa mitovirus 1 (NC_040543; *Mitoviridae*). ORFs translated with the mitochondrial genetic code are marked a mitochondria icon. For clarity, some lineages were collapsed (a non-collapsed version of the tree is available as Supplementary Information).

While the closest homologs of these narna-like viruses were identified in fungi, oomycete (protist) and marine arthropod samples, all three samples that contain these viruses are Bacillariophyta species (*A. radiata* and *P. pungens*) (Table 1, Figure 5). As their genome sequences share ~12% pairwise identity with other *Narnaviridae* we propose that Amphitrite narna-like virus, Poseidon narna-like virus and Halia narna-like virus represent novel species within the genus *Narnavirus*.

#### 3.4.2 *Mitoviridae*-like sequences

Seven RdRp protein sequences, retrieved from diverse algae host lineages – Rhodophyta, Haptophyta, Chlorophyta and Bacillariophyta – are related to members of the *Mitoviridae* (Figure 5). According to their placement in the *Mitoviridae* phylogeny as well as their level of divergence to existing mitoviruses (Figure 5, Table 1), these seven new viruses are potential members of the genus *Mitovirus*. All these mitovirus-like sequences have similar genome organizations, with the exception of one putative mitovirus with a genome that seemingly encodes a single RdRp-containing ORF (Figure 5). It is also notable that the RdRp-encoding ORFs from Aiolos mito-like virus, Asopus mito-like virus and Daimones mito-like virus can only be predicted using the mitochondrial code (Figure 5).

#### 3.4.3 *Leviviridae*-like sequences

One viral RdRp-like hit, in the Chlorophyta species *Pycnococcus provasolii,* is related to some bacteria-infecting *Leviviridae* and based on the levels of sequence identity this likely constitutes a new genus in this family (Table 1). As there were some bacterial reads in the *Pycnococcus provasolii* samples (MMETSP1471) (Figure 3B), it is likely that this Triton levi-like virus sequence infects bacteria (Actinobacteria or Proteobacteria-like) also present in the culture rather than *Pycnococcus provasolii*.

#### 3.4.4 *Picornavirales*-like sequences

Three sequences – denoted Pelias marna-like virus, Neleus marna-like virus and Tyro marna-like virus – were identified in diverse cultures belonging to various taxa (Figure 4): *Symbiodinium sp.* (Dinophyceae), *V. litorea* (Xanthophyceae) and *T. antarctica* (Bacillariophyta). These viruses exhibit sequence similarity with ssRNA+ viruses from the order *Picornavirales*. Specifically, they fell within the large algal associated family *Marnaviridae* (Figure 2C) and based on their respective positions in the phylogeny and the level of sequence divergence, Pelias marna-like virus could constitute a new genus in the *Marnaviridae*, while Neleus marna-like virus and Tyro marna-like virus are likely members of the genera *Kusarnavirus* and *Sogarnavirus*, respectively (Figure 6, Table 1). They also seem to share similar genome lengths and organizations as their closest relatives (Figure 6).

**Figure 6.**
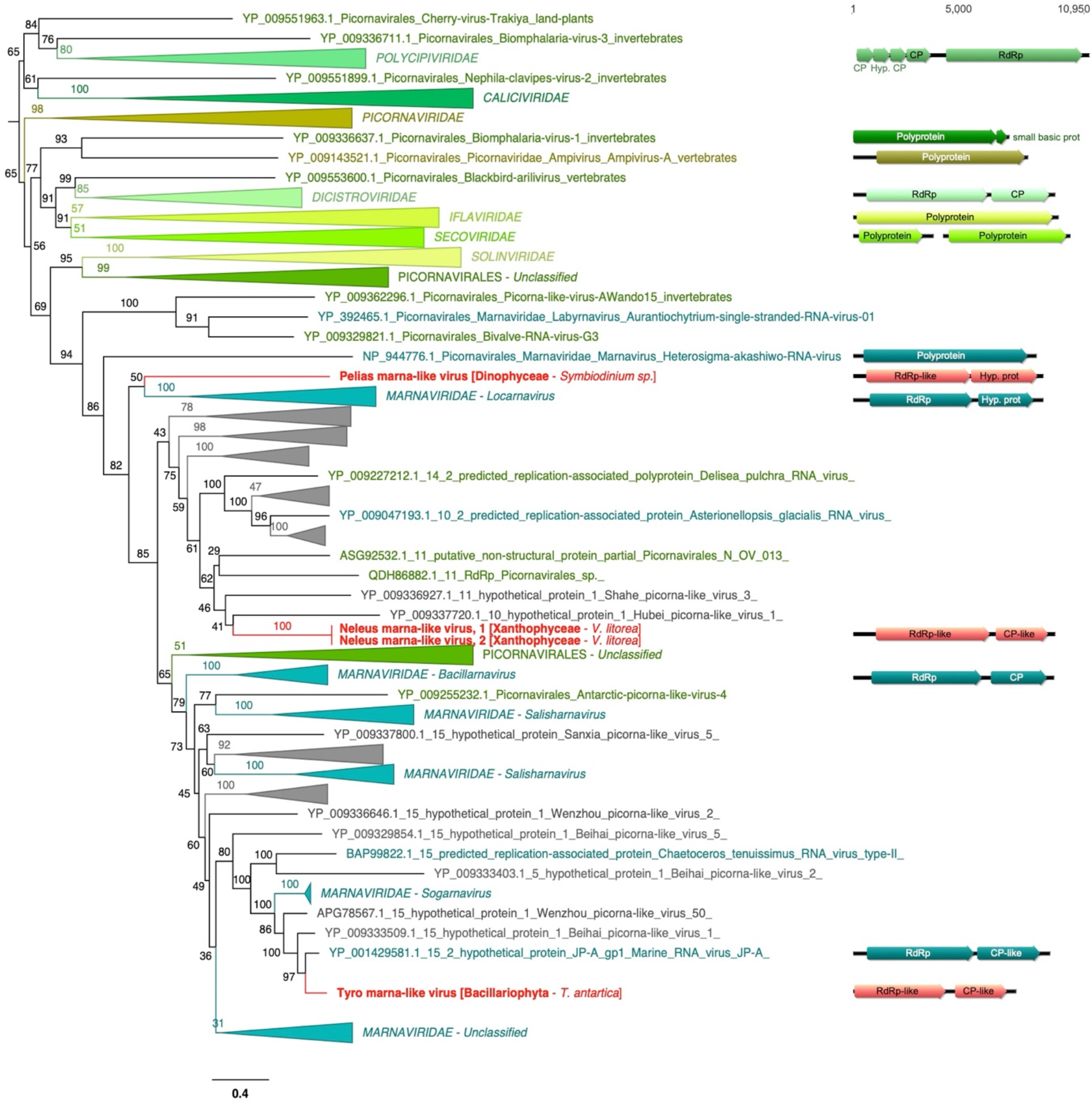
Phylogenetic placement of the newly described RNA virus sequences in the order *Picornavirales*. Left, ML phylogeny of the *Picornaviruses* RdRp (assuming the LG+F+R10 amino acid substitution model). Newly described viruses are indicated in red. Algae host taxon and species are specified in brackets. Branch labels = bootstrap support (%). The tree is mid-point rooted for clarity only. Right, genomic organisation of newly described viruses (red), closest homologs and the following *Picornavirales* order RefSeq representatives: Solenopsis invicta virus 2 (NC_039236; *Polycipiviridae*), Porcine enteric sapovirus (NC_000940; *Caliciviridae*), Foot-and-mouth disease virus - type O (NC_039210; *Picornaviridae*), Acute bee paralysis virus (NC_002548; *Dicistroviridae*), Infectious flacherie virus (NC_003781; *Iflaviridae*), Cowpea severe mosaic virus (NC_003544/NC_003545; *Secoviridae*). For clarity, some lineages were collapsed (a non-collapsed version of the tree is available as Supplementary Information).

### 3.5 Double-stranded (dsRNA) viruses

Almost a third of the RNA viruses newly reported here were related to dsRNA viruses of the family *Totiviridae* (Figure 2D). The single exception was the more divergent Charybdis toti-like virus, the exact placement of which within the order *Ghabrivirales* was unclear as it occupied a basal position in the phylogenetic tree and showed only low levels of sequence similarity to related viruses (~30% at RdRp protein level) (Figure 7, Table 1).

**Figure 7.**
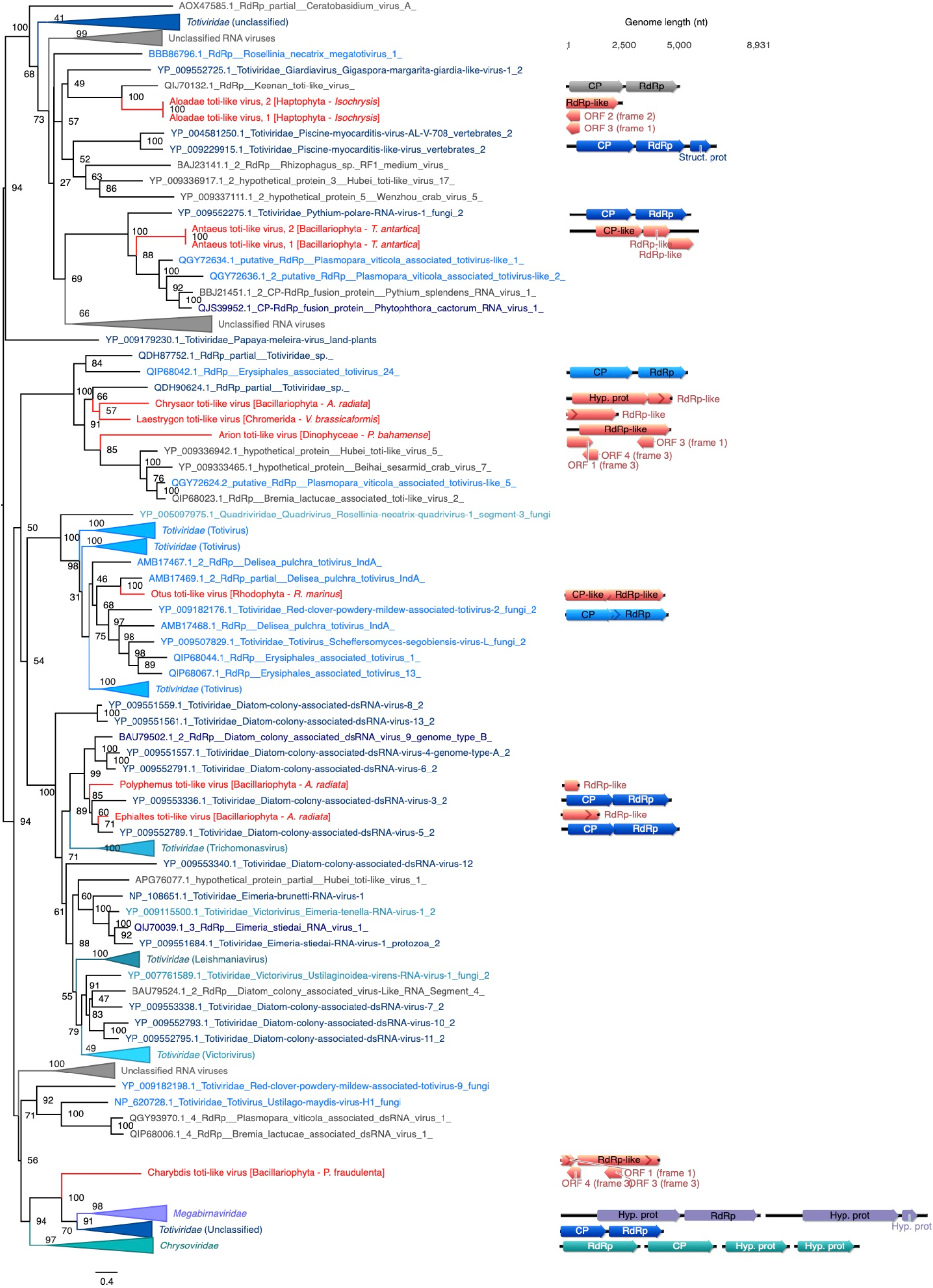
Phylogenetic position of the newly described RNA virus sequences among the *Ghabrivirales*. Left, ML phylogeny of the *Ghabrivirales* RdRp (assuming the LG+F+R10 amino acid substitution model). Newly described viruses are indicated in red. Algae host taxon and species are specified in brackets. Branch labels = bootstrap support (%). The tree is mid-point rooted for clarity only. Right, genomic organisation of the newly described viruses (red), closest homologs and the following representative *Ghabrivirales*: Rosellinia necatrix megabirnavirus 1/W779 (NC_013462/NC_013463; *Megabirnaviridae*), Tuber aestivum virus 1 (NC_038698; *Totiviridae*), Penicillium chrysogenum virus (NC_007539/NC_007540/NC_007541/NC_007542; *Chrysovirida*e). For clarity, some lineages were collapsed (a non-collapsed version of the tree is available as Supplementary Material).

Aloadae toti-like virus, found in Haptophyta *Isochrysis sp*, groups with the protist-associated *Giardiavirus* genus of the *Totiviridae*, and more surprisingly with Keenan toti-like virus recently identified in ectoparasitic flies (Figure 7), although with very high levels of sequence divergence (Table 1). Similarly, Chrysaor toti-like virus, Laestrygon toti-like virus and Arion toti-like virus, retrieved from Bacillariophyta, Chromerid and Dinophyceae, respectively, form a clade with *Totiviridae*-like sequences identified in either marine arthropods or oomycete protists (Figure 7). While these likely constitute a newly genus within the *Totiviridae*, their host remains uncertain. Antaeus toti-like virus, retrieved from the Bacillariophyta *T. antarctica,* groups with Pythium polare RNA virus 1 that infects the oomycete *Pythium polare*, confirming the presence of a polar stramenopile clade in the *Totiviridae*. Otus toti-like virus, identified in the Rhodophyta *R. marinus*, clusters (51% sequence identity) with the *Delisea pulchra totivirus* identified in the Rhodophyta (Figure 7).

Two additional toti-like viruses – Polyphemus toti-like virus and Ephialtes toti-like virus – were identified in *A. radiata* (Bacillariophyta) and, together with the diatom colony associated dsRNA viruses, form a new dsRNA viral clade, and likely genus, specifically associated with Bacillariophyta (diatoms) (Figure 7).

Strong similarities in genome organization were observed between the Otus toti-like virus and Antaeus toti-like virus and their toti-like homologs, with a potential single segment encoding a coat protein (CP) in 5’ and a RdRp in 3’ (Figure 7). As Charybdis toti-like virus, Chrysaor toti-like virus, Laestrygon toti-like virus, Arion toti-like virus, Polyphemus toti-like virus and Ephialtes toti-like virus all had partial genomes we were unable to determine their genomic organization, aside from the observation that they are likely unsegmented as they fall within the unsegmented *Totiviridae*. Unfortunately, such an assumption cannot be made for Charybdis toti-like virus, because of its basal position within the *Ghabrivirales*.

We identified six RdRp hits to members of the *Durnavirales* order of dsRNA virus (Figure 2C). With the exception of Aethusa amalga-like virus and Aegean partiti-like virus, their exact position within the six families that comprise this order (*Partitiviridae*, *Hypoviridae*, *Picobirnaviridae* and *Amalgaviridae*) is unclear due to their basal phylogenetic position (Figure 8). Moreover, these sequences seemingly have no association with specific microalgal groups, being observed in species of Rhodophyta, Bolidophyceae, Bacillariophyta, Chlorophyta and Dinophyceae (Figure 4). Aethusa amalga-like virus, retrieved from the Rhodophyta *R. marinus*, is clearly related to the *Amalgaviridae* (Figure 2 and Figure 8) and displays a moderate level of sequence divergence (43% identity in the RdRp) with *Zygosaccharomyces bailii virus Z* identified in fungi (Table 1). Whether this constitutes a new genus within the *Amalgaviridae* remains to be determined.

**Figure 8.**
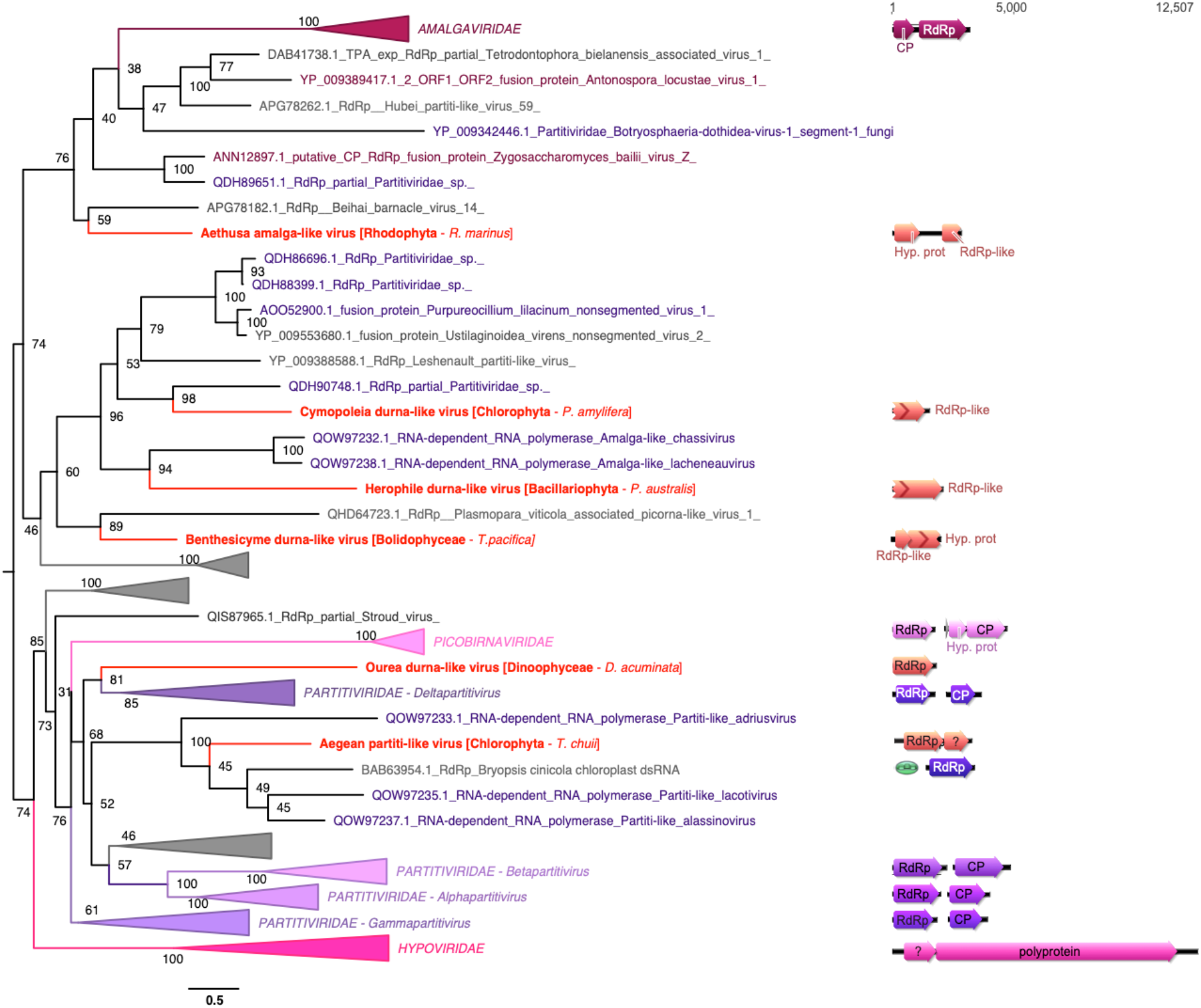
Phylogenetic positions of the newly described RNA viruses among the *Durnavirales*. Left, ML phylogeny of the *Durnavirales* RdRp (assuming the LG+F+R8 amino acid substitution model). Newly described viruses are indicated in red. Algae host taxon and species are specified in brackets. Branch labels = bootstrap support (%). The trees are mid-point rooted for clarity only. Right, genomic organisation of newly-discovered viruses (red), closest homologs and the following Partiti-picobirna super-clade representatives: Zygosaccharomyces bailii virus Z (NC_003874; *Amalgaviridae*), Cryphonectria hypovirus 2 (NC_003534; *Hypoviridae*), Chicken picornavirus (NC_003534/ NC_040438; *Picobirnaviridae*), Fig cryptic virus (NC_015494/NC_015495; *Deltapartitivirus*), Discula destructiva virus 1 (NC_002797/NC_002800; *Gammapartitivirus*), Ceratocystis resinifera virus 1 (NC_010755/NC_010754; *Betapartitivirus*), White clover cryptic virus 1 (NC_006275/NC_006276; *Alphapartitivirus*). ORFs translated with the plastid genetic code are labelled with a green plastid. For clarity, some lineages were collapsed (a non-collapsed version of the tree is available as Supplementary Information).

Three other viruses, Benthesicyme durna-like virus, Herophile durna-like virus and Cymopoleia durna-like virus, were related to the Amalga-like lacheneauvirus and Amalga-like chassivirus, both previously identified in cultures of *Ostreobium* sp. (Chlorophyta), and that fell between the *Amalgaviridae* and *Partitiviridae* families in our phylogenetic analysis (Figure 8). The genomic sequences for Benthesicyme durna-like virus, Herophile durna-like virus and Cymopoleia durna-like virus were likely partial such that their organization, particularly whether they comprise one of two segments, could not be established (Figure 8).

Aegean partiti-like virus falls in the *Partitiviridae*, grouping with the Partiti-like lacotivirus, Partiti-like allasinovirus, Partiti-like Adriusvirus and Bryopsis cinicola chloroplast dsRNA (BDRC): these are all *Partitiviridae* and associated with Ulvophyceae algae (Figure 8). The presence of Aegean partiti-like virus in *Tetraselmis chuii* (Chlorophyta) strongly supports the existence of a Chlorophyta-infecting partiti-like viral genus. Assuming a homologous genome organization, the genome of Aegean partiti-like virus would comprise a single segment encoding a RdRp in its 5’ region as well as a hypothetical protein, potentially a coat protein, in the 3’ region. Whether Aegean partiti-like virus is associated with the host chloroplast remains uncertain. Finally, Ourea durna-like virus is highly divergent and falls basal to the bi-segmented *Partitiviridae* (Figure 8). However, considering the length and the single ORF organization of the partial genomic sequence retrieved, it is likely that a second segment encoding a CP may not have been detected by BLAST due to very high levels of sequence divergence.

### Negative-sense viruses (ssRNA-)

A novel RdRp sequence, Susy yue-like virus, was identified in the *Pseudo-nitzschia heimii* (Bacillariophyta) culture. This virus clusters among the ssRNA-*Haploviricotina*, falling between the *Qinviridae* and the *Yueviridae* families (Figure 9). Considering the length of the RdRp segment and the bi-segmented genome organization of related members of the *Qinviridae* and *Yueviridae* (Figure 9), it is highly likely that the Susy yue-like virus genome is partial. In similar manner to the *Qinviridae*, Susy yue-like virus has an IDD sequence motif instead of the common GDD triad in the catalytic core of its RNA virus replicase (RdRp), although the functional implications of this alternative motif are unclear.

**Figure 9.**
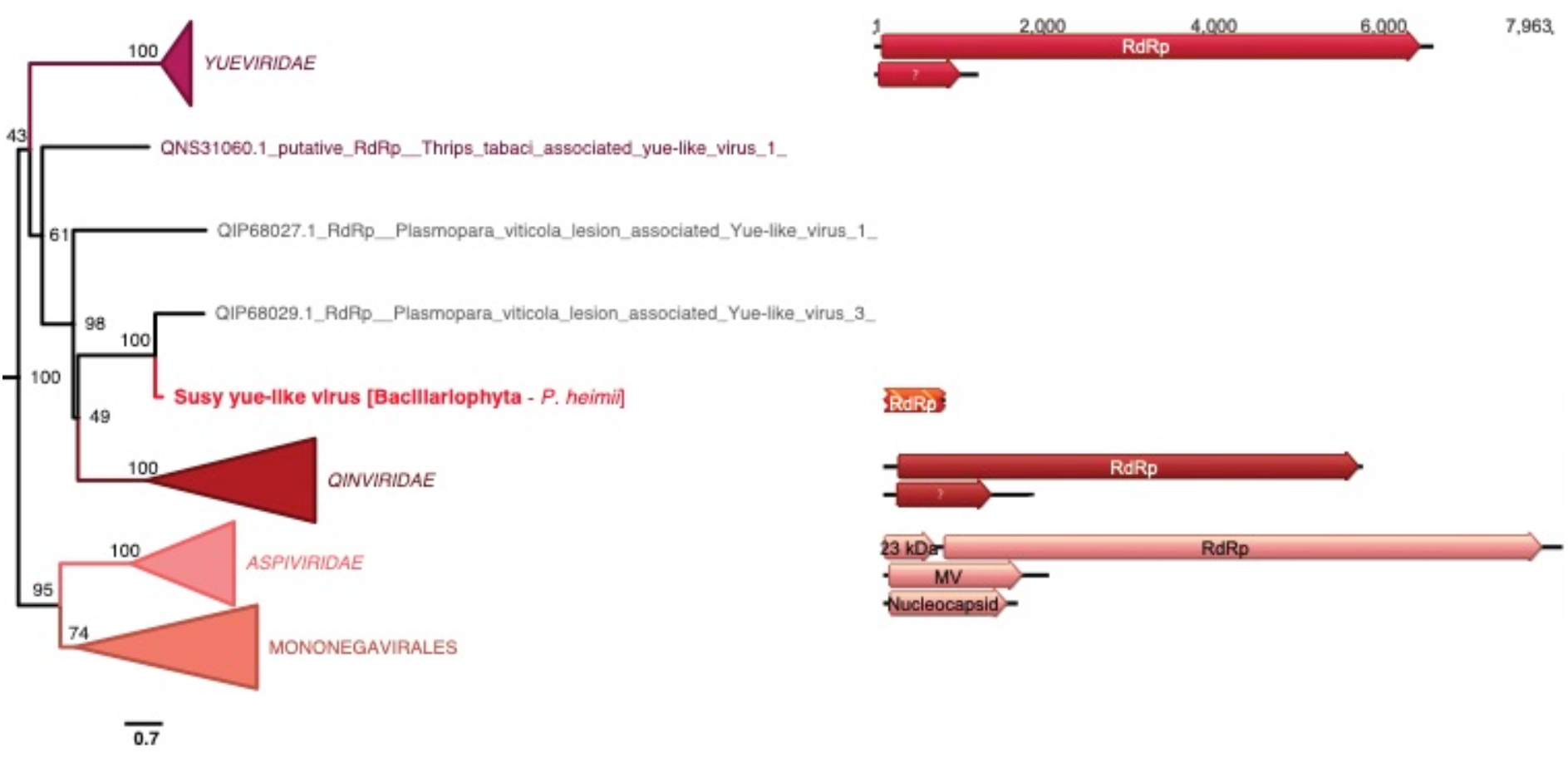
Position of the newly described RNA virus in the phylum *Haploviricotina.* Left, ML phylogeny of the *Haploviricotinia* RdRp (employing the LG+F+R10 amino acid substitution model). The virus newly described here is shown in red. Algae host taxon and species are specified in brackets. Branch labels = bootstrap support (%). The tree is mid-point rooted for clarity only. Right, genomic organisation of the newly described virus (red) and the following homologs representatives: Shahe yuevirus-like virus 1 (NC_033289/NC_033290; *Yueviridae*), Beihai sesarmid crab virus 4 (NC_032274/NC_032272; *Qinviridae*), Blueberry mosaic associated virus (NC_033754/NC_036634/NC_036635; *Aspiviridae*). For clarity, some lineages were collapsed (a non-collapsed version of the tree is available as Supplementary Information).

### Detection of divergent RNA viruses based on RdRp motifs and structural features

The microalgal transcriptomes sequenced as part of the MMETSP likely contain viruses that are highly divergent in sequence, sharing only limited sequence similarity to those currently available and hence challenging to detect using BLAST-based methods. To identify RNA viruses at lower levels of homology, we conducted an extensive analysis utilising RdRp protein functional motifs and structural features on all the BLAST-unannotated sequences: this accounted for 10-34% of the total predicted ORFs of at least 200 amino acid residues in length (Figure S2).

A very large proportion of the sequences retained from our combined RdRp-based HMM, InterproScan analysis were false-positive hits as they were either confidently detected as eukaryotic-like sequences using Phyre2 or were too distant to be safely considered as an RdRp (i.e. unreliable alignment and no detection of RdRp catalytic motifs) (Table S3). However, five RdRp-like candidates were retained from the manual curation steps. While no robust RdRp-like signal could be detected using Phyre2 (i.e. prediction confidence scores below 90%) (Table S3), the presence of a significant HMM-detected homology with the PROSITE PS50507 profile (i.e. RdRp of ssRNA+ virus catalytic domain profile; Table S2) enabled us to further analyze these candidates as potential RdRp sequences.

Four of these five RdRps came from the genus *Bigelowiella*, and three (MMETSP0045_DN12861, MMETSP1054_DN18666 and MMETSP1052_DN19445) shared high identity levels (>90% at both protein and nucleotide levels), while MMETSP1359_DN14104 shared only 70% identity (Table 2). Although the PROSITE PS50507 profiles were built from ssRNA+ RdRp sequences, the IDD C-motif exhibited by these four RdRp-like candidates is found in the ssRNA- *Qinviridae*-like viruses as well as the new Susy yue-like virus found in *Pseudo-nitzschia heimii* (MMETSP1423). However, the nucleotide sequences of these RdRp-encoding contig candidates exhibited a strong match (e-value < 1E-90) with a genome contig (BIGNAscaffold_41_Cont1731) from the *Bigelowiella natans* genome (GCA_000320545.1). Hence, rather than representing an exogenous RNA virus, the distant RdRp hit in this case most likely constitutes an endogenous viral element (EVEs) indicative of a past, and likely ancient, infection event.

**Table 2.** RdRp-like hits retrieved from the HMM-profile and Phyre2 analyses. Presence of the A, B and C motifs are noted along with the sequence of the C-motif.

| Contig ID | Taxon | RdRp profile | E-value | A | B | C | Phyre2 confid % | %ID | Hit info |
| --- | --- | --- | --- | --- | --- | --- | --- | --- | --- |
| MMETSP1359_DN14104<br>_c0_g1_i1_len843_1 | *Bigelowiella longifila*<br>(Cercozoa) | PS50507 | 6.0E-07 | Yes | ? | IDD | 64.2 | 16 | PDB header: transferase |
| MMETSP0045_DN12861<br>_c0_g1_i1_len664_1 | *Bigelowiella natans*<br>(Cercozoa) | PS50507 | 8.7E-06 | Yes | ? | IDD | 40.7 | 24 | DNA/RNA polymerases |
| MMETSP1054_DN18666<br>_c0_g1_i1_len657_1 | *Bigelowiella natans*<br>(Cercozoa) | PS50507 | 8.9E-06 | Yes | ? | IDD | 41.6 | 24 | DNA/RNA polymerases |
| MMETSP1052_DN19445<br>_c0_g1_i1_len738_1 | *Bigelowiella natans*<br>(Cercozoa) | PS50507 | 1.0E-05 | Yes | ? | IDD | 40.4 | 24 | DNA/RNA polymerases |
| MMETSP0202_DN4292<br>_c0_g1_i1_len814_1 | *Karenia brevis*<br>(Dinophyceae) | PS50507 | 4.6E-05 | Yes | ? | GDT | 56.7 | 17 | PDB header: hydrolase |

In the case of the remote RdRp-like signal in MMETSP0202_DN4292, no GDT sequence at motif C could be identified in an expansive RdRp data set^35^. Hence, it is unclear if MMETSP0202_DN4292 is a true viral RdRp or a false-positive hit.

## 4. Discussion

To the best of our knowledge we report the largest survey of RNA viruses in microalgal curated cultures. With the discovery of 30 new and divergent viruses, 29 of which are likely to infect algae species in which no viruses have previously been reported, this study greatly extends our knowledge of the microalgae RNA virosphere. More broadly, this work demonstrates the potential of protists to be major reservoirs of novel RNA viruses.

Despite the viral diversity documented, only 6% (33 of 570) of the transcriptomes analysed here contained evidence of an RNA virus, far lower than equivalent meta-transcriptomic studies of single organisms^36–38^. The use of purified cultures is expected to reduce the number of viruses compared to direct environmental samples, preventing the sequencing of co-circulating viruses as well as those infecting other microorganisms in the environment. However, this relative paucity of RNA viruses could also reflect methodological limitations. First, the lack of rRNA depletion in the library processing leads a concomitant reduction in the number of non-rRNA transcripts, including those from viruses. Indeed, most of the viruses reported here display very low transcript abundance, suggesting that additional RNA viruses may have been undetected due to poor sequencing coverage. Second, the limited number of viruses identified is likely to reflect the high levels of sequence divergence expected for protist viruses compared to those currently available in sequence databases. Indeed, many of the viruses identified in this study share less than 30-40% sequence identity, toward what might be the limit of a viable BLAST-based analysis. Hence, this study has been conducted at the boundaries of the detectable virosphere, with a myriad of more divergent viruses yet to be discovered.

### 4.1 RNA virus are widespread among lineages of unicellular algae

Our knowledge of RNA viruses associated with microalgae is scarce. The small number reported so far are mostly associated with a specific subset of algal species from the Bacillariophyta and Chlorophyta, ignoring the wide diversity of microalgae (Figure 1A). We extend this diversity by revealing, for the first time, RNA viruses (i.e. RdRp sequences) in the Haptophyta, Chromeraceae (Alveolates), as well as in the Stramenopiles Xanthophyceae and Bolidophyceae. We also identified new virus-algae clade associations. For example, we present the first observation of *Picornavirales*, *Ghabrivirales* (*Totiviridae*) and *Durnavirales* (*Partititivridae*) in Dinophyceae cultures, *Lenarviricota* and *Durnavirales* in Rhodophyta cultures, and *Durnavirales* in Bacillariophyta cultures. Importantly, our study also constitutes the first observation of a *Muvirales*-like ssRNA-virus in a Bacillariophyta sample, perhaps only the second negative-sense RNA virus identified in microalgae.

With the exception of *Symbiodinium* sp. for which a ssRNA+ virus was previously reported^39,40^, all the viruses described in this study represent the first observation of an RNA virus in each respective host species. In addition, none of the 73 microalgal viruses reported previously were identified here. More generally, the distribution of RNA viruses obtained in this study, comprising ssRNA+, ssRNA- and dsRNA viruses, varies considerably between taxa and likely reflects sampling bias rather than a host specificity of RNA virus infection. These factors might have contributed to the lack of viral identification in poorly investigated and divergent taxa such as Euglena, Glaucophytes and Cryptophytes. Further studies with particular emphasis on these taxa are clearly required.

The first observation of an ssRNA- virus in a Bacillariophyta, together with the previous observation of a bunya-like virus reported in the distantly-related Chloroarachniophyte *C. reptans* (Cercozoa) and bunya-like siRNAs in brown algae (Phaeophyta)^41^, again demonstrates that microalgae can be infected with negative-sense RNA viruses. Interestingly, the related *Qinviridae* and *Yueviridae* have been exclusively identified from meta-transcriptomic studies conducted on marine arthropods holobionts, such that algae could constitute the true hosts for most of these viruses^42,43^. Undoubtedly, the presence of ssRNA-viruses in microbial eukaryotes needs to be further characterized.

### 4.2 *Narnaviridae*-like and *Mitoviridae*-like viruses are common in microalgal cultures

A third of the viruses reported here were from the order *Lenarviricota* that includes the *Narnaviridae* and *Mitoviridae* and often characterised by a single RdRp ORF^44^. Although they were initially thought to be restricted to fungi, these seemingly simple RNA viruses appear to be more widespread than initially thought. Indeed, *Narnaviridae*-like viruses have recently been associated with a wide range of protist organisms, including protozoan parasites like *Plasmodium vivax*^45–48^ and the oomycete *Phytophthora infestans*^49^, while narna-like viruses have also been detected in diatoms^50^. Similarly, the *Mitoviridae* were considered as exclusively infecting fungi, until the recent discovery of the Chenopodium quinoa mitovirus 1 in a plant^51^ and mito-like viruses in the Chlorophyta *Osteobium* sp.^52^ led their host range to be re-evaluated. The three new narna-like viruses in Bacillariophyta discovered here, as well as the proposal of seven new mitovirus-like species in algal lineages as diverse as Haptophyta, Bacillariophyta, Rhodophyta and Chlorophyta, provides further evidence for the ubiquity of these viruses in protists.

Whether all the mitoviruses documented here are associated with the mitochondria, as typical of the *Mitoviridae*, remains to be determined. In addition, while the unique RdRp-encoding segment has already been demonstrated as sufficient for virus infectivity, recent studies have suggested the presence of an additional segment, without an assigned function, in both *Leptomonas seymouri* and *Plasmodium vivax*^45,48^. Whether the viruses newly described here have unsegmented or bipartite genomes remains to be determined. Most of the *Lenarviricota*-like sequences described here display ambigrammatic ORFs, with their reverse strand encoding additional ORFs. This feature has already been reported in narnaviruses and could represent a potential solution to extreme genome compaction^53–55^.

The growing evidence for the extended host range of both *Narnaviridae* and *Mitoviridae* beyond the fungal clades has important consequences in our knowledge of the early events in the evolution of eukaryotic RNA viruses. Indeed, the ubiquity of *Mitoviridae* and *Narnaviridae* in eukaryotes is compatible with the protoeukaryotic origins of these viruses and the bacterial *Leviviridae*, such that they are relics of a past endosymbiont infection of a eukaryotic ancestor. Accordingly, cytoplasmic *Narnaviridae* would have escaped from mitochondria to the more RNA hospitable cytosol^3^. In addition, *Narnaviridae* and *Mitoviridae* are not associated with cellular membranes^56^, which could also reflect their ancient origin from a protoeukaryote ancestor without cellular compartments.

### 4.3 The extension of the *Marnaviridae* to new algal taxa

Most of the algal RNA viruses described to date belong to the order *Picornavirales*^8^, including the *Marnaviridae*. Currently, the *Marnaviridae* comprise 20 species, distributed among seven genera based on their capsid similarities. Notably, all these viral species are associated with marine samples or algae cultures^57^. The three picorna-like viruses newly identified in this study fell within the *Marnaviridae*. Despite similar genome organizations, these three viruses have relatively high levels of divergence from known *Marnaviridae*, in turn suggesting that the *Marnaviridae* diversity has only been sparsely sampled. This diversity will very likely increase with the sequencing of phytoplankton cells. While the detection of Neleus marna-like virus and Tyro marna-like virus in Bacillariophyta and Xanthophyceae could reflect the specificity of *Sogarnavirus* and *Kusarnavirus* to Stramenopile algae, the first detection of a *Marnaviridae*-like virus in the Dinophyceae species *Symbiodinium* sp. suggests that the host range of this algal-infecting viral family is not restricted to Stramenopile eukaryotes.

### 4.4 The ancestry of the *Durnavirale*s and *Ghabrivirales* dsRNA viruses

Approximately half of the RNA viruses identified in this study are related to the *Totiviridae* (*Ghabrivirales*) and *Partitiviridae* (*Durnavirales*) families of dsRNA virus. The *Totiviridae* currently comprises 28 formally-assigned species divided into five genera^32,58^. Interestingly, *Totiviridae* are exclusively associated with unicellular eukaryotes, with two of the five *Totiviridae* genera associated with latent fungal infections (*Totivirus* and *Victorivirus*), while *Trichomonasvirus, Giardiavirus* and *Leishmaniavirus* have been associated with protozoan parasite infections^32^.

Each of the new *Totiviridae*-like sequences identified here were retrieved from a range of algal hosts spread among diverse branches of the microbial eukaryote tree (Bacillariophyta, Dinophyceae, Haptophyceae, Rhodophyta and Chromeraceae). Hence, as with the *Marnaviridae*, the diversity of the *Totiviridae* has likely been greatly underestimated. In addition, some of the novel viruses identified cluster with totiviruses previously reported in Bacillariophyta diatoms^59,60^ and the Rhodophyta *Delisea pulchra*^61^. These observations support the existence of a Bacillariophyta and a Rhodophyta-infecting clade in the genus *Totivirus* that will need to be confirmed with studies of additional species. It was also notable that other toti-like viruses identified here cluster with viruses found in non-algal hosts, such as invertebrates (ticks, crustaceans), fungi and protozoan parasites. While host mis-annotations cannot be formally excluded, the presence of *Totiviridae* in protozoan parasites, fungi and algae could signify that the host range of the *Totiviridae* is far larger than appreciated.

Six dsRNA-like new viruses identified here show clear homology with those of the order *Durnavirales*, including the *Partitiviridae* and the *Amalgaviridae* that comprise bi-segmented and unsegmented dsRNA viruses, respectively. The *Partitiviridae* are classified into five genera and mainly associated with plants and fungi, although more recently with oomycetes^62^ and to Apicomplexa^63^. The *Amalgaviridae* comprise two genera associated with either fungi (*Zybavirus* genus) or land plants (*Amalgavirus* genus)^58,64^. In addition to the recent association of newly described partiti- and amalgavirus-like viruses in the microalgae *Ostreobium* sp. (Cholorophyta)^52^, our identification of these novel and divergent *Durnavirales*-like viruses in several distant algae taxa again suggests that host range for this viral order has been underestimated.

### 4.5 Are cryptic viruses a common feature of unicellular eukaryotes?

RNA viruses causing host cell lysis and hence mortality are commonly reported^65^, with an emblematic example being the lysis of the harmful algal bloom-forming diatoms, haptophytes and dinoflagellates, leading to bloom collapse^66,67^. Although we did not aim to assess the phenotypic effects of viral infection on algal hosts, it is noticeable that most of the viruses identified here were related to the *Totiviridae*, *Partitiviridae*, *Mitoviridae* and *Narnaviridae*, all previously reported as associated with cryptic and persistent infections^32^. This is consistent with the design of the MMETSP study that would tend to identify non-pathogenic viruses. It is also in accordance with the growing evidence that a non-neglectable component of RNA virus-host associations are symptomless or even beneficial to their host, with potentially importations evolutionary implications^68,69^.

### 4.6 Limitations to virus discovery and inferring virus-host relationships

A key element of this study was use of mono-strain cultures, which were axenic whenever possible, enabling more accurate virus-host assignments. While Bacteria, and to a lesser extent, Archaea, were present in the non-axenic cultures, the placement of most of the newly described viruses within eukaryotic-infecting viral families clearly supports their association with algae. Despite this, some of the newly- described viruses were associated with viral lineages traditionally associated with fungal or metazoan hosts. This likely reflects the lack of representation of microalgal viruses in current sequence databases or a mis-annotation to secondary metazoan host, particularly given the recent efforts to describe the fungal virome^70–73^. Similarly, many of the newly identified viruses share homology with viruses identified in metagenomics studies on marine invertebrates^36^. It is widely established that such similarities to holobiont virome studies should be treated with caution, as the viruses reported could in fact be infecting symbionts, eukaryotic parasites, or bacteria that are also present in these samples^3^. Marine invertebrate organisms are also important ocean filters and virus removers^74^, again compatible with the idea that at least some of the viruses identified here may infect other marine organisms.

We also attempted to identify more distant RNA viruses using a protein profile and structural-based approach. However, no remote RNA virus signals could be confidently detected using this method, although a distant endogenous viral element in *Bigelowiella* was identified. While the *de novo* prediction of protein 3D structures has experienced major improvements over the last decade^75^, revealing robust homology strongly relies on structural comparisons and modelling based on pre-existing structures^22^. Critically, however, only a very limited number of non-human viruses are available among the viral proteins deposited in the Protein Data Bank. This poor representativeness of protein structures is a major roadblock in the ability to detect highly divergent RdRps. Indeed, a better characterization of RdRp structures combined with the enrichment of RdRp motif and profile databases will help counter the challenge posed by the high levels of sequence divergence in protist samples and the concomitant loss of detectable evolutionary signals. In addition, the high percentage of false positives in the HMM analysis highlights the need to increase and optimize the sensitivity and stringency of such methods.

While our study significantly extends our knowledge of RNA virus diversity among unicellular eukaryotes, experimental confirmation is needed to formally assign such viruses to their specific microalgae hosts and to assess the impact of viral infection on host biology. Perhaps more importantly, additional effort is needed to detect the signal of remote sequence homology in the highly divergent RNA viruses that are likely commonplace in protists.

## Acknowledgments

SM thanks the Moore Foundation for funding her involvement in the MMETSP project. ECH is funded by an Australian Research Council Australian Laureate Fellowship (FL170100022).

## Data availability

All viral genomes and corresponding sequences detected in this study will be deposited in the NCBI GenBank and SRA upon the acceptance. The accessions ID will be listed in Table 1.

