## Supplementary Information for "Revealing RNA virus diversity and evolution in unicellular algae transcriptomes"

**Figure S1.** Comparison of RNA virus and algae host *rbcL* transcript abundance. *rbcL* gene accessions (NCBI GenBank) are provided in Table S4.

**Figure S2.** Ratio of annotated versus unannotated ORFs in each algal clade. The percentage of ORFs (>200 amino acids) with entries in nr using Blastp (e-value e-05) and in profile databases using InterProScan (defaut cut-off) are indicated in green and yellow, respectively. The proportion of ORFs (>200 amino acids) for which no entries are reported shown indicated in grey.

**Table S1.** MMETSP samples used in this study.

**Table S2.** List of PFAM and PROSITE RdRp-profiles used in the HMM-based detection.

**Table S3.** HMM-profile and Phyre2-based detection results. All RdRp-like hits obtained using HMMer3 were also submitted to Phyre2. Presence or absence of the RdRp canonical motifs A, B or C and alternative C-motifs are also indicated.

**Table S4.** Relative abundance of *RbcL* host gene sequences in RNA virus-positive libraries. *rbcL* abundances correspond to the number of reads per million.

**Table S5.** Additional information for MMETSP samples that are positive for RNA viruses.

**Tree (Newick) files:** Non-collapsed versions of phylogenetic trees shown in Figures 5-9.

- Lenarviricota_FULL-TREE.newick
- Picornavirales_FULL-TREE.newick
- Ghabrivirales_FULL-TREE.newick
- Durnavirales_FULL-TREE.newick
- Haploviricotina_FULL-TREE.newick
