## Supplementary Table 2 for "Revealing RNA virus diversity and evolution in unicellular algae transcriptomes"

**Table S2.** List of PFAM and PROSITE RdRp-profiles used in the HMM-based virus detection.

| **Database** | **Name** | **Description** |
| --- | --- | --- |
| PFAM | PF05919 | Mitovirus RdRp |
| PFAM | PF04197 | Birnavirus RdRp |
| PFAM | PF17501 | Viral RdRp (C-terminal domain) |
| PFAM | PF00680 | Viral RdRp (RdRP_1) |
| PFAM | PF00978 | RdRP_2 |
| PFAM | PF00998 | Viral RdRp (RdRP_3) |
| PFAM | PF02123 | RdRP_4 |
| PFAM | PF07925 | RdRP_5 |
| PFAM | PF04196 | Bunyavirus RdRp |
| PFAM | PF00946 | *Mononegavirales* RdRp |
| PFAM | PF05788 | Orbivirus RdRp (VP1) |
| PFAM | PF08467 | Luteovirus P1-P2 |
| PFAM | PF00603 | Influenza RdRp (subunit PA) |
| PFAM | PF12289 | Rotavirus RdRp (VP1) |
| PFAM | PF00604 | Influenza RdRp (subunit PB2) |
| PFAM | PF00602 | Influenza RdRp (subunit PB1) |
| PFAM | PF00972 | Flavivirus RdRp |
| PFAM | PF14314 | *Mononegavirales* RdRp (Methyltranferase region) |
| PFAM | PF12426 | Viral RdRp |
| PFAM | PF06478 | Coronavirus RdRp (N-terminal) |
| PFAM | PF06317 | Arenavirus RdRp |
| PFAM | PF03431 | *Leviviridae* RdRp (beta-chain) |
| PROSITE | PS50524 | *Birnaviridae* RdRp (catalytic domain) |
| PROSITE | PS50523 | *Reoviridae* RdRp (catalytic domain) |
| PROSITE | PS50522 | RNA-containing bacteriophages RdRp (catalytic domain) |
| PROSITE | PS50526 | *Mononegavirales* (non-segmented) RdRp (catalytic domain) |
| PROSITE | PS50525 | *Mononegavirales* (segmented) RdRp (catalytic domain) |
| PROSITE | PS50507 | ss(+)RNA virus RdRp (catalytic domain) |
