## Supplementary Table 4 for "Revealing RNA virus diversity and evolution in unicellular algae transcriptomes"

**Table S4.** Relative abundance of *RbcL* host gene sequences in RNA virus-positive libraries. *rbcL* abundances correspond to the number of reads per million.

| **SRA acc** | **rbcL gene acc.** | **Species** |
| --- | --- | --- |
| **SRR1294395** | EF520333.1 | *Pseudo-nitzschia fraudulenta* |
| **SRR1294402** | MN808826.1 | *Rhodosorus marinus* |
| **SRR1296718** | NC_038008.1 - rbcL | *Astrosyne radiata* |
| **SRR1296745** | EF520333.1 | *Pseudo-nitzschia fraudulenta* |
| **SRR1296746** | EF520333.1 | *Pseudo-nitzschia fraudulenta* |
| **SRR1296747** | EF520333.1 | *Pseudo-nitzschia fraudulenta* |
| **SRR1296822** | HQ661109.1 | *Rhodella maculata* |
| **SRR1296825** | FJ002140.1 | *Amphiprora sp* |
| **SRR1296826** | FJ002140.1 | *Amphiprora sp* |
| **SRR1296875** | HF931099.1 | *Tetraselmis chuii* |
| **SRR1296920** | DQ514795.1 | *Thalassiothrix antarctica* |
| **SRR1296921** | DQ514795.1 | *Thalassiothrix antarctica* |
| **SRR1296939** | MK642517.1 | *Chaetoceros curvisetus* |
| **SRR1296959** | DQ514795.1 | *Thalassiosira antarctica* |
| **SRR1296962** | DQ514795.1 | *Thalassiosira antarctica* |
| **SRR1296965** | DQ514795.1 | *Thalassiosira antarctica* |
| **SRR1300363** | AF372696.1 | *Triparma pacifica* |
| **SRR1300394** | FM207547.1 | *Pseudo-nitzschia pungens* |
| **SRR1300423** | AY119783.1 | *Isochrysis CCMP1244* |
| **SRR1300433** | AF298221.1 | *Symbiodinium sp.* |
| **SRR1300439** | AY119783.1 | *Isochrysis CCMP1244* |
| **SRR1300541** | U30280.1 | *Pycnococcus provasolii* |
